# A novel thermostable aspartic protease from *Talaromyces leycettanus* and its specific autocatalytic activation through an intermediate transition state

**DOI:** 10.1101/528265

**Authors:** Yujie Guo, Tao Tu, Yaxin Ren, Yaru Wang, Yingguo Bai, Xiaoyun Su, Yuan Wang, Bin Yao, Huoqing Huang, Huiying Luo

## Abstract

Aspartic proteases exhibit optimum enzyme activity under acidic condition and have been extensively used in food, fermentation and leather industries. In this study, a novel aspartic protease precursor (pro*Tl*APA1) from *Talaromyces leycettanus* was identified and successfully expressed in *Pichia pastoris*. Subsequently, the auto-activation processing of the zymogen pro*Tl*APA1 was studied by SDS-PAGE and N-terminal sequencing, under different processing conditions. *Tl*APA1 shared the highest identity of 70.3 % with the aspartic endopeptidase from *Byssochlamys spectabilis* (GAD91729) and was classified into a new subgroup of the aspartic protease A1 family, based on evolutionary analysis. Mature *Tl*APA1 protein displayed an optimal activity at 60 °C and remained stable at temperatures of 55 °C and below, indicating the thermostable nature of *Tl*APA1 aspartic protease. During the auto-activation processing of pro*Tl*APA1, a 45 kDa intermediate was identified that divided the processing mechanism into two steps: formation of intermediates, and activation of the mature protein (*Tl*APA1). The former step was completely induced by pH of the buffer, while the latter process depended on protease activity. The discovery of the novel aspartic protease *Tl*APA1 and study of its activation process will contribute to a better understanding of the mechanism of aspartic proteases auto-activation.

**IMPORTANCE:** The novel aspartic protease *Tl*APA1 was identified from *T. leycettanus* and expressed as a zymogen (pro*Tl*APA1) in *P. pastoris*. Enzymatic characteristics of the mature protein were studied and the specific pattern of zymogen conversion was described. The auto-activation processing of pro*Tl*APA1 proceeded in two stages and an intermediate was identified in this process. These results describe a new subgroup of aspartic protease A1 family and provide insights into a novel mode of activation processing in aspartic proteases.

## INTRODUCTION

Proteases (EC 3.4.11-24) make up a large share of the total global industrial enzymes (1). They have extensive applications in the dairy, baking, beverages, brewing, meat and functional food industries (1, 2). On the basis of the optimal pH of hydrolysis, proteases have been grouped into three categories–acidic, neutral, and alkaline proteases. They can also be classified into serine, cysteine, metallo and aspartic proteases, depending upon their catalytic residues. Aspartic proteases (EC 3.4.23) have two aspartic residues at their catalytic center, which are vital for hydrolytic cleavage of peptide bonds (3). The activity of aspartic proteases can be specifically inhibited by pepstatin A. Molecular weights of aspartic proteases commonly range between 30 to 50 kDa, while some can weigh up to 55 kDa. Aspartic proteases are generally considered as acidic proteases, because they have isoelectric points of 3.0–4.5 and show optimal activity at pH 3.0–5.0. Most aspartic proteases have an optimal temperature in the range of 30–50 °C, while some exhibit maximum activity at 55 °C. Most aspartic proteases are also sensitive to high temperatures and show poor thermostability, limiting their applications to mesophilic conditions. Thus, the thermal stability of aspartic proteases has been the subject of attention in many recent studies (4). The discovery of novel thermostable enzymes, especially from extremophiles, is a potential method to tackle the aforesaid problem (2, 5).

Aspartic proteases are wide spread in many organisms–vertebrates, insects, plants, fungi and even viruses have been widely reported as sources of aspartic proteases (6). The production of aspartic proteases from fungi has several advantages including short productive cycle, simple late purification, and low costs (7). Most commercial aspartic proteases used currently in industrial production are derived from filamentous fungi. Aspartic proteases from fungi are mainly categorized into two groups–pepsin-like and rennin-like enzymes (8). The pepsin-like enzymes include aspergillopepsin (9), penicillopepsin (10), trichodermapepsin (11) and rhizopuspepsin (12); while the rennin-like enzymes are mainly produced by *Mucor*, *Rhizomucor* and *Chryphonectria* (3). However, the yield of fungal aspartic proteases by industrial fermentation is usually low. An aspartic protease from *Aspergillus foetidus* was extracellularly produced with a activity of only 63.7 U/mL (13). *Pichia pastoris* is an excellent expression system that has been effectively used to solve the problems of low yield of proteases. Many aspartic proteases have been heterogeneously expressed in *P. pastoris* (14, 15, 16). An aspartic protease from *Rhizomucor miehei* was produced in *P. pastoris* with the activity of 3480.4 U/mL (17).

Typical aspartic proteases are initially synthesized in the form of inactive precursors (zymogens), which protect host cells from proteolysis (6). The functions of the N-terminal pro-segments of aspartic proteases have been studied extensively and include facilitating correct folding, blocking the active site, and stabilizing the protein (18, 19). It is generally accepted that the propeptides are auto-catalytically cleaved at acidic pH (20, 21, 22), and their further processing by other peptidases is important for the activation of aspartic proteases from *Candida parapsilosis* (23). Crystal structures of some aspartic protease zymogens and their activation intermediates have been reported (20, 21, 24, 25, 26), which contribute to the understanding of propeptide interactions with catalytic proteins. Current studies on zymogen activation have mainly concentrated on aspartic proteases associated with diseases (22, 27), yet little is known about this process among aspartic proteases from fungi. There is evidence that the predicted zymogens vary in length depending on each fungus, suggesting their unique activation processes (6). Therefore, studying the activation processing of fungal aspartic proteases is of great significance to understand the mode of activation of the whole family.

Given the importance of novel and thermostable aspartic proteases in industrial processes, a gene coding for a novel thermostable aspartic protease, *Tlapa1*, was found and cloned from the thermophilic filamentous fungus *Talaromyces leycettanus*. Phylogenetic analysis indicated that *Tl*APA1 belonged to a new subgroup of aspartic proteases A1 family. Moreover, *Tl*APA1 has been previously expressed in *P. pastoris* in its zymogen form and its auto-activation has been studied in detail. The auto-activation process of *Tl*APA1 was affected by pH and enzymatic activity, and occurred in two stages distinguished by the presence of a processing intermediate. The mature proteases with activity were subsequently purified and characterized biochemically. In this study, a novel thermostable aspartic protease was discovered and synthesized as a zymogen in *P. pastoris*, and its autocatalytic activation was studied.

## RESULTS

### Gene cloning and sequence analysis

The gene encoding a novel aspartic protease zymogen was identified from the genome of *Talaromyces leycettanus* and named as pro*Tlapa1*. The DNA sequence (MK108371 in the GenBank database) had 1275 bp consisting of 3 introns and 4 exons. pro*Tlapa1* encode a polypeptide of 424 amino acids including a putative signal peptide of 19 residues and a propeptide of 61 residues at the N-terminus. The molecular mass and p*I* value were estimatedto be 42.5 kDa and 4.7, respectively. Three *N*-glycosylation sites (N144, N253 and N357) were predicted using the NetNGlyc 1.0 Server. The deduced amino acid sequence of pro*Tlapa1* shared the highest identity of 70.3 % with the aspergillopepsin A-like aspartic endopeptidase from *Byssochlamys spectabilis* (GAD91729, 28), and of only 41.4% with a functionally characterized aspergillopepsin-1 from *Aspergillus oryzae* RIB40 (Q06902, 29). Among aspartic proteases with determined three-dimensional structures, pro*Tlapa1* (residues 86 to 405) showed the highest identity (41.7%) with the corresponding domain of the mature aspartic protease (PDB 1IZD) from *Aspergillus oryzae* (30).

### Phylogenetic analysis of *Tl*APA1

The results of BLASTP analysis showed that *Tl*APA1 belonged to the A1 family of aspartic proteases. However, this family of proteases comprises of many subgroups with complex evolutionary relationships. To obtain a clear evolutionary relationship between *Tl*APA1 and other homologs of the A1 family, a phylogenetic analysis based on the amino acid sequence alignment was performed using MEGA 7.0. These results indicated that aspartic proteases from different microorganism were separated from each other in the evolutionary tree (Fig. 1). Seven subgroups that had been reported in previous studies emerged in the process of evolution in the following order: aspergillopepsin, penicillopepsin, trichoderpepsin, podosporapepsin, endothiapepsin, rhizopuspepsin, and murcorpepsin subgroups. However, *Tl*APA1 did not belong to any of these subgroups. *Tl*APA1 along with Q4WZS3 from *Aspergillus fumigates* belonged to a new clade in the evolutionary tree (Fig. 1). This clade emerged after the formation of the rhizopuspepsin subgroup in the evolutionary tree. The evolutionary position of *Tl*APA1 suggested that *Tl*APA1 might have unique characteristics distinct from other aspartic peptidases.

**Fig. 1.**
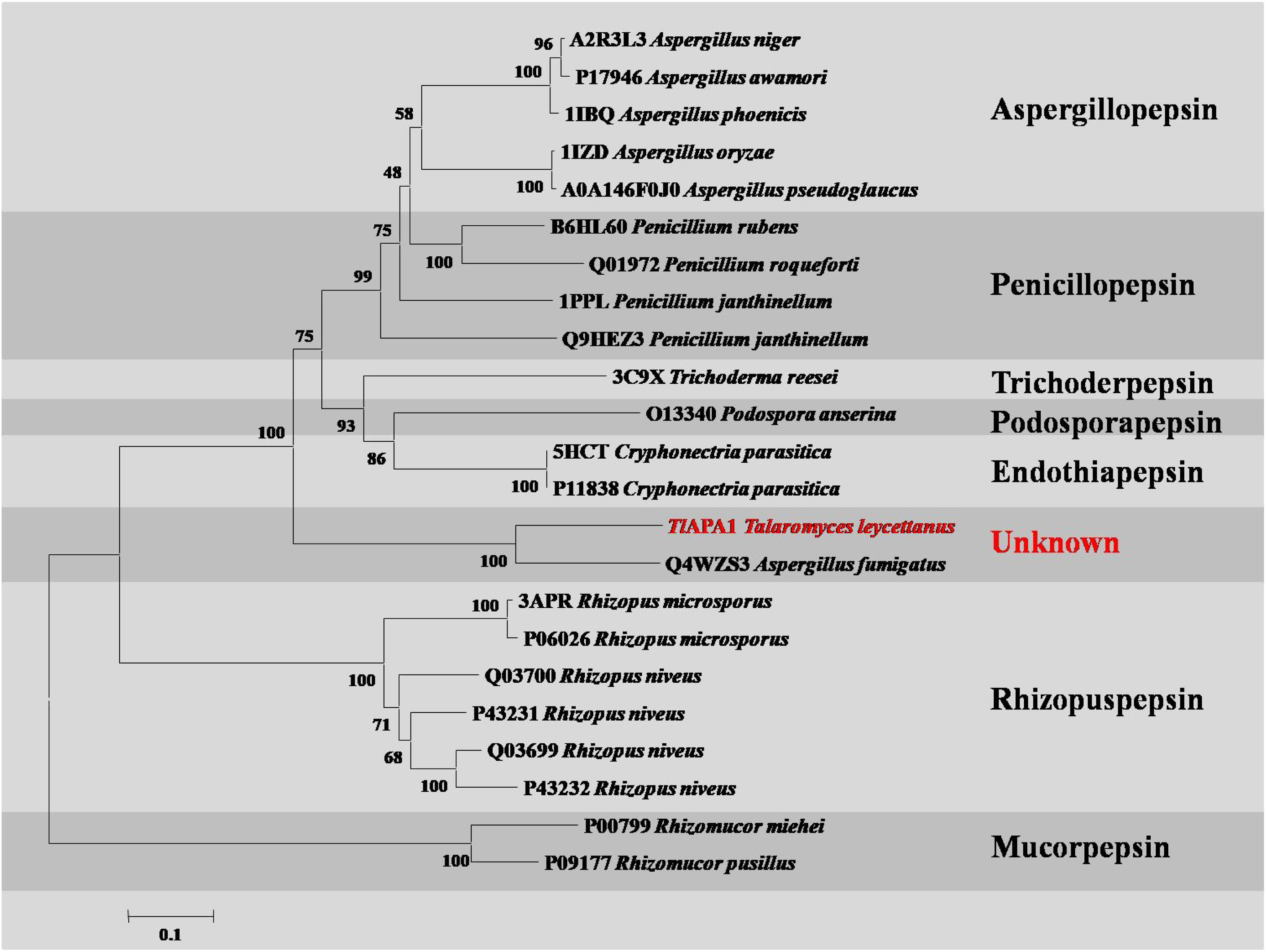
Phylogenetic analysis of the subgroups of aspartic proteases A1 family. The amino acid sequences of A1 family were obtained by a BLAST analysis using the *Tl*APA1 protein (GenBank accession number MK108371) as the query sequence. The evolutionary tree was constructed by the neighbor-joining method. The sequences are labeled with their GenBank accession numbers and host fungi. Numbers indicated in the tree branches are the bootstrap values (%) based on 1000 replications. The subgroups of A1 family are classified using gray shadow and names are indicated in over striking. *Tl*APA1 is shown in red which, together with Q4WZS3, belonged to a new unknown clade.

### Heterologous expression and purification of proenzyme

The zymogen pro*Tl*APA1, consisting of the N-terminal propeptide and the mature domain, was expressed in *P. pastoris* GS115. Expression of the recombinant protein was confirmed by electrophoresis. As shown by SDS-PAGE (Fig. 2), a specific protein band corresponding to a molecular mass of approximately 53 kDa was obtained, which was higher than the calculated value (45 kDa) of pro*Tl*APA1. Upon treating the samples with Endo H, the target band appeared at approximately 45 kDa. The first five residues of purified pro*Tl*APA1 were V-P-A-P-S, as identified by N-terminal amino acid sequence analysis. These results indicated that *Tl*APA1 was produced in the form of a zymogen in *P. pastoris* GS115.

**Fig. 2.**
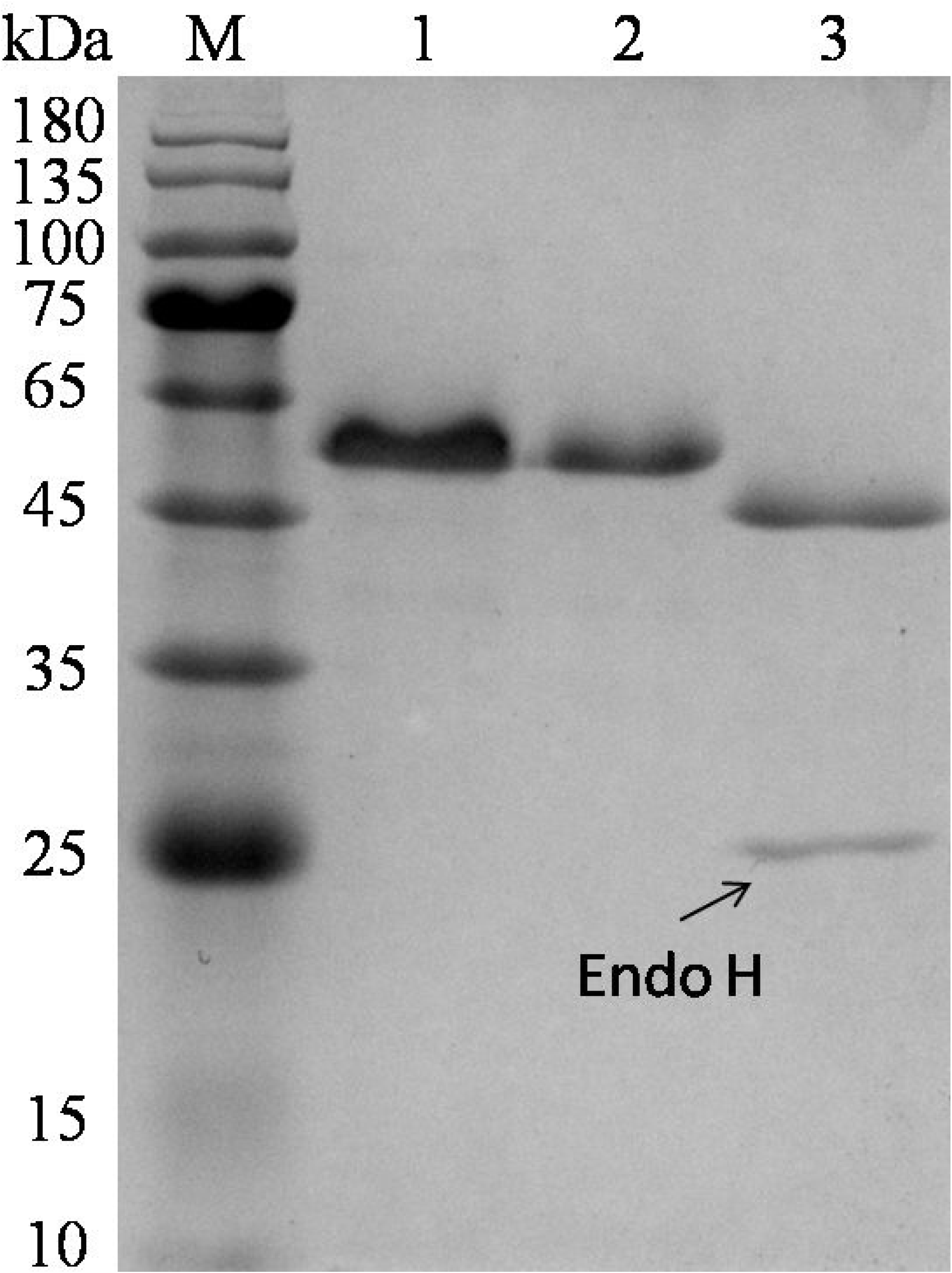
SDS-PAGE analysis of the recombinant pro*Tl*APA1. Lane M, molecular mass standard; Lane 1, crude zymogen pro*Tl*APA1; Lane 2, purified recombinant pro*Tl*APA1; and Lane 3, deglycosylated zymogen pro*Tl*APA1 treated with Endo H.

### Process of auto-activation

The inactive precursors of aspartic proteases were usually auto-catalytically activated under acidic conditions. In this study, the processing of the zymogen conversion was determined by SDS-PAGE and N-terminal amino acid sequencing. As shown in Fig. 3, the activation process had been already initiated at 0 min of incubation at a room temperature (about 25 °C), indicating the rapidity of the process, potentially due to disintegration of zymogens in the reaction whose pH had been adjusted to 3.5 before incubation. The appearance of two new bands was accompanied by the weakening of the zymogen band (50 kDa) before 30 min of incubation (Fig. 3). The molecular weights of the two new products were 45 kDa and 40 kDa, and their first 5 residues were identified as L-D-F-E-P and V-A-Q-P-A, respectively. After 90 min of incubation at room temperature at pH 3.5, the pro*Tl*APA1 was completely converted to a 40 kDa band (Fig. 3). We suspected that the 45 kDa product was an intermediate in the conversion process of pro*Tl*APA1 to mature *Tl*APA1, and the processing sites on this 45 kDa product were confirmed by N-terminal sequencing. The above results indicated that the precursors pro*Tl*APA1 could be processed into its mature form in an intermolecular manner, and two processing sites (L67-L68, D85-V86) of pro*Tl*APA1 auto-activation were identified.

**Fig. 3.**
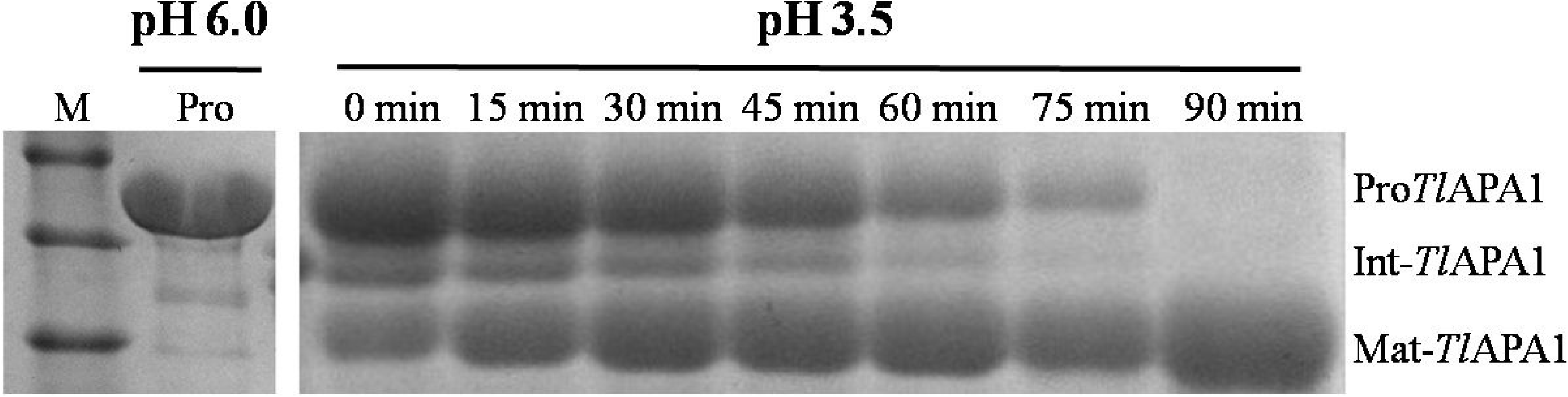
Analysis of pro*Tl*APA1 auto-activation for 90 min. The time course of processing at room temperature (∼20 °C) was analyzed using 12% SDS-PAGE. Pro*Tl*APA1, recombinant precursor without the signal peptide; Int-*Tl*APA1, intermediate produced during auto-activation processing; Mat-*Tl*APA1, mature protein after auto-activation.

### Effects of proteolytic activity on auto-activation processing

As previous studies have illustrated, processing induced auto-activation was related to its own proteolytic activity. Pepstatin A is a specific inhibitor of aspartic protease, which can effectively inhibit its protease activity. Hence, the effect of pepstatin A on the zymogen conversion of *Tl*APA1 was examined at a concentration of 5 μM of pepstatin A. Electrophoretic analyses revealed that upon pepstatin A treatment, the apparent molecular mass of the pro*Tl*APA1 decreased from 50 to 45 kDa (Fig. 4). The first 5 amino acid residues of processed proteins (45 kDa) were determined as L-D-F-E-P, which were identical to the cleavage intermediate of *Tl*APA1. This result indicated that auto-processing intermediates (45 kDa products) were produced in the presence of pepstatin A. However, the mature protein (40 kDa) was not obtained even upon prolonged the incubation time up to 3 h. This demonstrated that the activation process from intermediates to mature proteins was inhibited by pepstatin A. Hence, we concluded that the activation process from intermediates to mature protein was accompanied by proteolytic processing. We also studied the effect of catalytic residue of *Tl*APA1 on zymogen conversion by replacing the catalytic Asp103 residue with Asn. This showed that although the catalytic mutant did not prevent processing of the precursor into the 45 kDa intermediate, it terminated the auto-activation of pro*Tl*APA1 in the intermediate stage (data not shown). The above results show that the two phases of the auto-activation process of pro*Tl*APA1 were independent of each other, and the activity of aspartic proteases had greater effect on the latter stage rather than on the former.

**Fig. 4.**
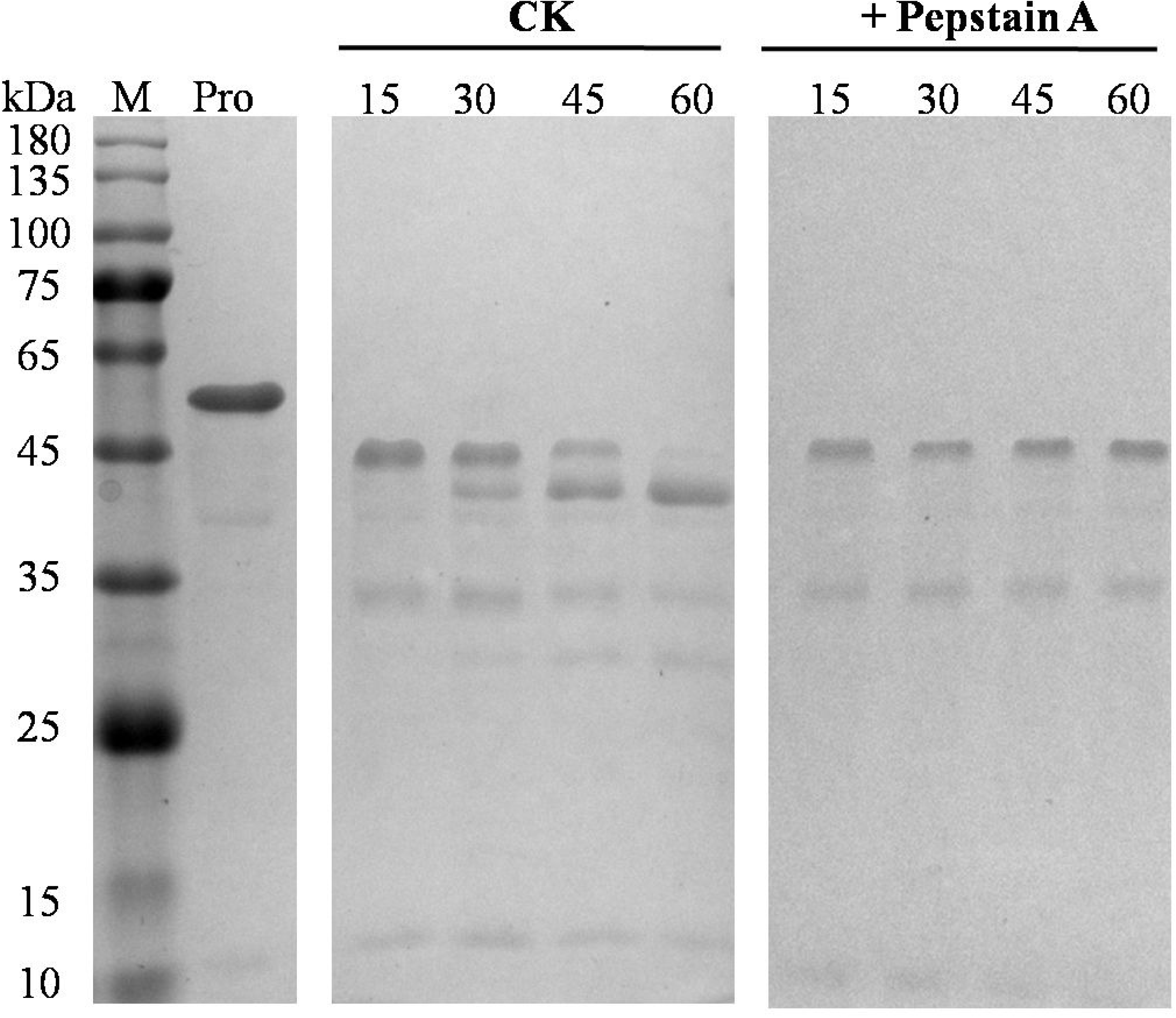
Effect of the inhibitor pepstain A on zymogen auto-processing. The auto-activation processing of recombinant zymogens with or without pepstatin A was analyze using SDS-PAGE at the indicated times. M, molecular mass standard; Pro, purified zymogens before auto-activation. The processing time of the auto-activation ranged from 15–60 (min).

### Effect of pH on auto-activation processing

The auto-activation processing of aspartic proteases is often triggered by low pH. To determine the optimum pH of zymogen conversion, this processing was executed in range of pH 3.0–6.0, and the proteolytic activities of treated samples were detected using casein (1%, w/v) as a substrate, during the processing of the aspartic protease zymogen. During prolonged incubation at pH 3.0, the proteolytic activities were increased and the enzyme activity peaked after 60 min (Fig. 5A). We speculated that the precursor was almost completely processed after 60 min of incubation at pH 3.0, which was confirmed by SDS-PAGE (Fig. 5B, pH 3.0). The processing induced auto-activation at pH 4.0 and pH 3.0 were comparable, although complete activation at pH 4.0 required longer incubation time (about 90 min). As shown in Fig. 5A, the increasing of proteolytic activities was absent at pH 6.0, and no differences in protein bands were observed (Fig. 5B, pH 6.0). This indicated that zymogen pro*Tl*APA1 was not converted at pH 6.0. Interestingly, the precursor band was cleaved at pH 5.0 (Fig. 5B), although the final activity was dramatically lower than at pH 3.0 and pH 4.0. In order to understand the reason for the lowered activity at pH 5.0, the N-terminal amino acid sequences of mature proteins at pH 3.0 and pH 5.0 were determined, and found to be V-A-Q-P-A and A-V-Q-G-G. This demonstrated that two different mature proteins were generated– *Tl*APA1 at pH 3.0 and M2-*Tl*APA1 at pH 5.0, however, the mature protein M2-*Tl*APA1 was inactive. In conclusion, precursors of aspartic proteases could be activated at the range of pH 3.0–4.0, while auto-activation occurred most efficiently at pH 3.0, which is closest to the optimum pH of mature aspartic protease.

**Fig. 5.**
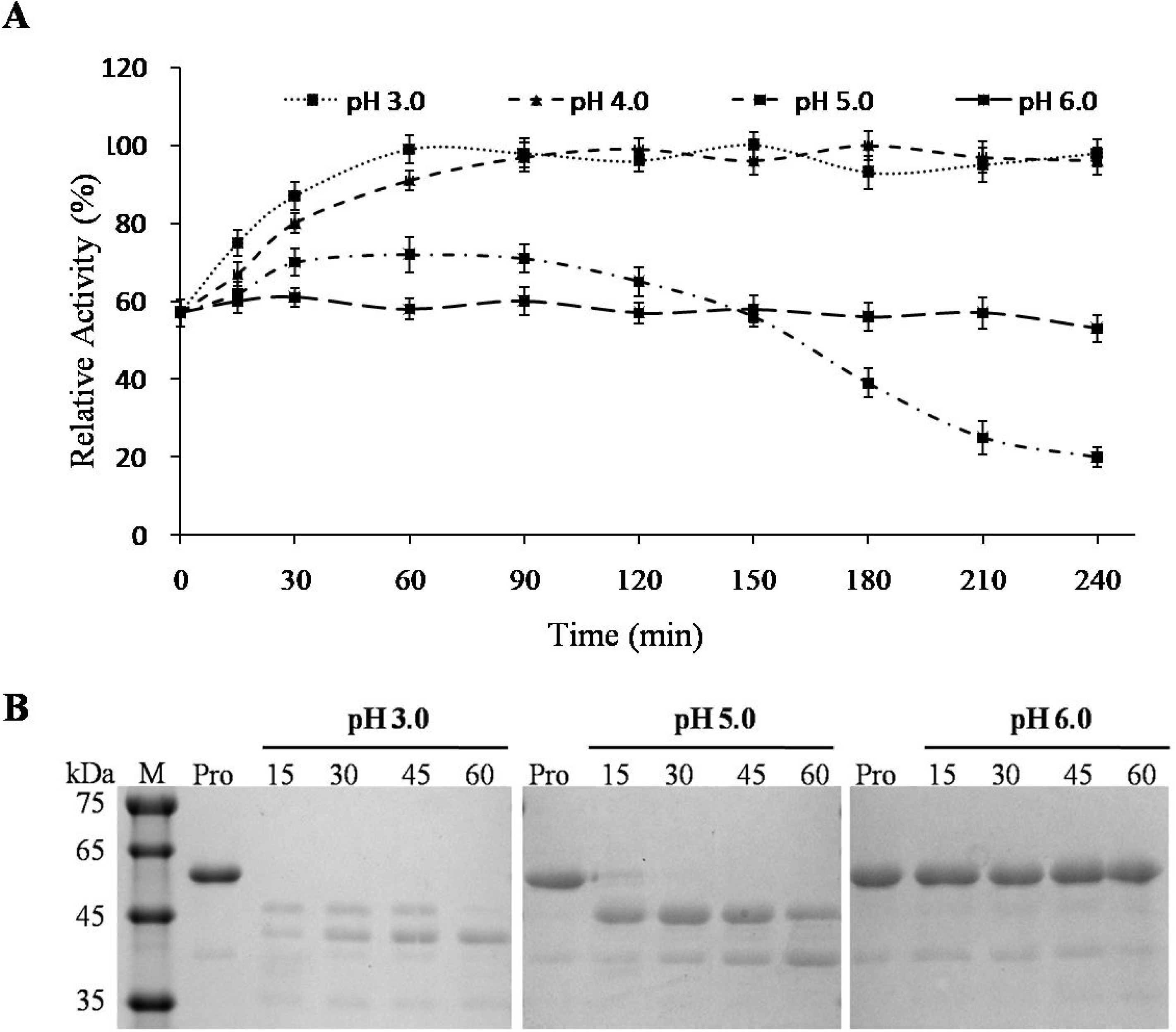
Analysis of pro*Tl*APA1 auto-activation at varying pH 3.0–6.0. (a) Proteolytic activities measured at different times with range of pH 3.0–6.0 at 60 °C. (b) SDS-PAGE analysis of auto-activation processing at 37 °C with pH 3.0, 5.0, and 6.0.

### Biochemical characterization of *Tl*APA1

The enzymatic characteristics of mature aspartic proteases *Tl*APA1 and M2-*Tl*APA1 were assessed using casein (1%, w/v) as a substrate. *Tl*APA1 showed the highest activity at pH 3.5 (Fig. 6A), similar to that that seen in most fungal aspartic proteases. As shown in Fig. 6B, *Tl*APA1 had an optimal temperature of 60 °C, which was higher than aspartic proteases obtained from most other fungi. We further measured stabilities of aspartic protease *Tl*APA1 under different pH and temperature conditions. *Tl*APA1 retained greater than 80 % of its initial activity after 60 min of incubation at 37 °C over a range of pH 2.0–6.0 (Fig. 6C). The stability at acidic pH makes *Tl*APA1 favorable for applications in food, beverages, and brewing industries. Fig. 6D shows that *Tl*APA1 was extremely stable below 55°C, retaining almost all of its initial activity after 1 h of incubation. At higher temperatures, half-life of *Tl*APA1 was 30 min at 60 °C and 5 min at 65 °C. The thermostability of *Tl*APA1 was higher than that of highly homologous aspartic proteases from other fungi. M2-*Tl*APA1 activity was not detected at any of the aforementioned temperature and pH conditions. Purified recombinant *Tl*APA1 had a specific activity of 2187.4 ± 67.3 U⋅mg^−1^, while the *K*_m_, *V*_max_, *k*_cat_, and *k*_cat_/*K*_m_ values were determined as 1.9 mg⋅mL^−1^, 2,321 μmoL⋅min^−1^⋅mg^−1^, 1,410 s^−1^ and 723.5 mL⋅s^−1^⋅mg^−1^, respectively.

**Fig. 6.**
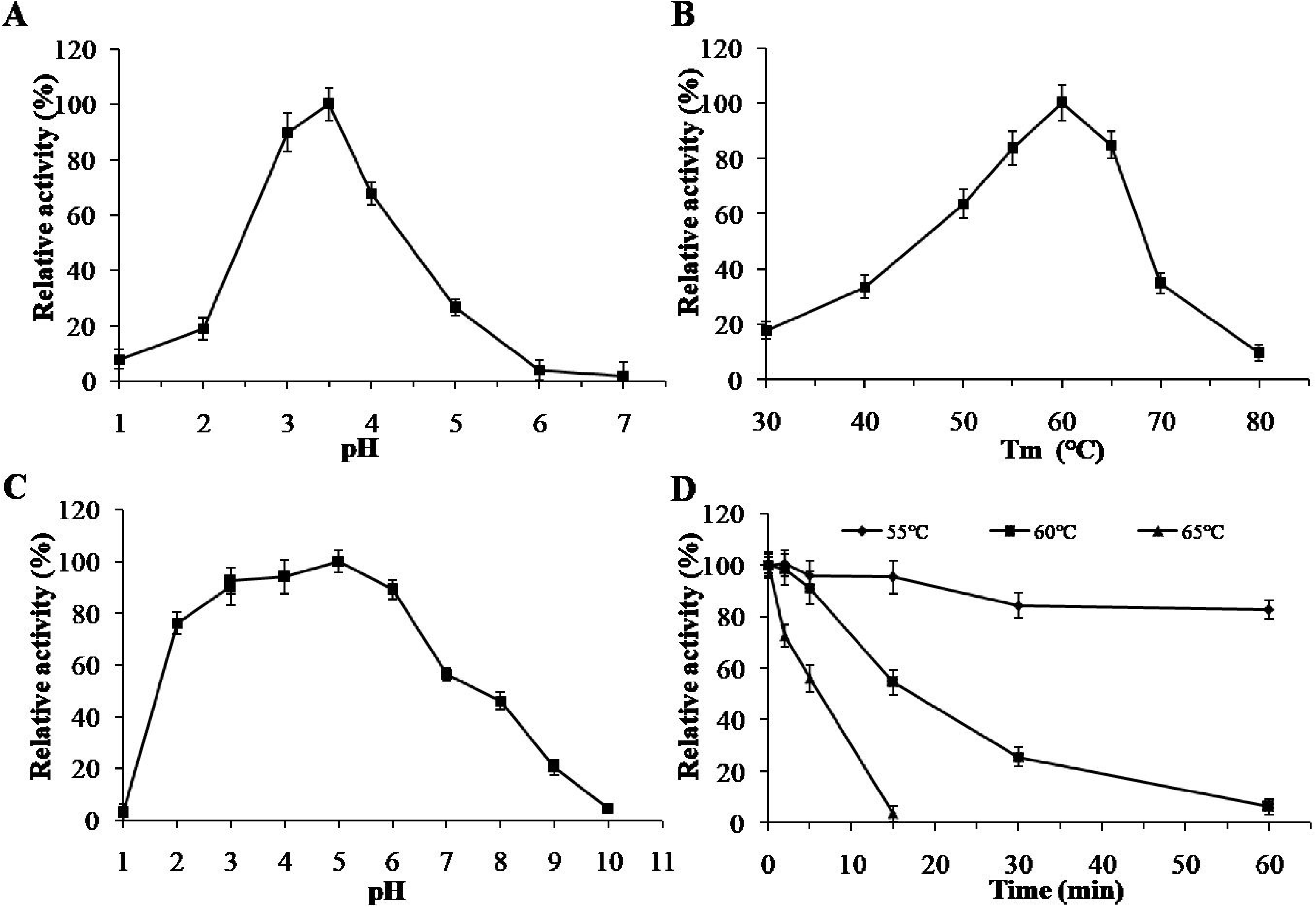
Characterization of purified mature enzyme *Tl*APA1. (a) pH-activity profile. (b) Temperature-activity profile. (c) pH-stability profile after 1 h-incubation at 37 °C at different pH values. (d) Temperature-stability profile after incubation at pH 3.0 and different temperatures for various durations.

## DISCUSSION

Novel enzymes with unique characteristics such as pH adaption, thermostability and tolerance to metal ions are in valuable for research and industrial purposes (31). Extremophiles are excellent sources of novel enzymes that can retain their integrity and function under extreme reaction conditions (32). *Talaromyces leycettanus* JCM12802, a thermotolerant fungus, that has an optimal growth temperature of 42 °C, is the source of various thermostable hydrolases including *β*-mannanase (33), xylanase (34), *β*-glucanase (35), and *α*-Amylase (36). In this study, a novel aspartic protease precursor comprising of 424 amino acid residues was identified in *Talaromyces leycettanus* and was determined to be a member of the A1 family of aspartic proteases.

A1 family, the most well studied one of the 16 families of aspartic proteases, is further subdivided into 5 super families (AA, AC, AD, AE, and AF) (37). Pfam protein family prediction indicated that *Tl*APA1 belonged to the A1 family, one that contains many biochemically-characterized enzymes including pepsin, chymosin, rennin, and cathepsin D (37, 38). When full sequences of the A1 family proteins were used for phylogenetic analysis, the phylogenetic tree indicated *Tl*APA1 as a sister to an uncharacterized aspartic protease Q4WZS3 from *Aspergillus fumigates*, whose clade has not yet been discovered (Fig. 1). Phylogenetic analysis is able to provide deeper insights into the evolution of a new clade. The phylogenetic tree showed that *Tl*APA1 largely differed with other homologs in terms of the amino acid sequence, indicating that *Tl*APA1 potentially had distinct enzyme characteristics and special applications like the mucorpepsin subgroup, a unique clade in the evolutionary process and an important class of proteases, widely used as milk coagulating agents (39). The evolution of A1 family has been well studied, which has contributed to an indepth understanding of fungal aspartic protease evolution (40). Fungal aspartic proteases have undergone large sequence diversification leading to their evolutionary complexity (37). To further investigate the evolutionary position of *Tl*APA1 among subgroups of A1 family, a phylogenetic analysis was performed and an evolutionary tree was constructed (Fig 1). Our analysis suggested *Tl*APA1 as a potential evolutionary intermediate linking rhizopuspepsins and other subgroups.

The full amino acid sequence of secreted aspartic proteases contains a propeptide region followed by a mature protein (6). The pro*Tl*APA1 zymogen was composed of an N-terminal propeptide of 61 residues, and a mature domain of 339 residues. Our results demonstrated that pro*Tl*APA1 could be processed auto-catalytically into the mature aspartic protease under acidic conditions, consistent with other homologs (41). However, the N-terminal propeptide of pro*Tl*APA1 precursor was processed auto-catalytically in two phases. Firstly, precursors were processed into intermediates driven by pH-dependent structural changes that were not affected by a specific inhibitor or the D103N mutation. Secondly, the process that turned intermediates into mature enzymes was auto-catalyzed by enzymes; that could be stalled in the presence of a specific inhibitor or mutation of active residues. We further studied the conversion processing of pro*Tl*APA1 at different pH conditions and determined that the optimum pH of conversion was similar to that of its peak activity. Interestingly, when we activated the zymogens at pH 5.0, a minor mature protein band was generated in SDS-PAGE. This auto-processing site (S98-A99) was also confirmed by N-terminal sequencing, and was located 13 residues downstream of the first mature site. However, the mature products at pH 5.0 had no proteolytic activity (Fig 7). The large differences in the processing of pro*Tl*APA1 compared to other homologs could arise from the N-terminal propeptide of *Tl*APA1 precursor (Fig.S1). The N-terminal propeptide of *Tl*APA1 precursor showed low sequence identity (< 30%) to other known aspartic proteases. The N-terminal propeptide of *Tl*APA1 precursor possessed an additional 61 residues with a greater abundance of arginine residues, compared to other homologs (Fig.S1). We thus speculated that the sequence peculiarity of N-terminal pro-segment lead to the special auto-activation processing of the *Tl*APA1 precursor.

**Fig. 7.**
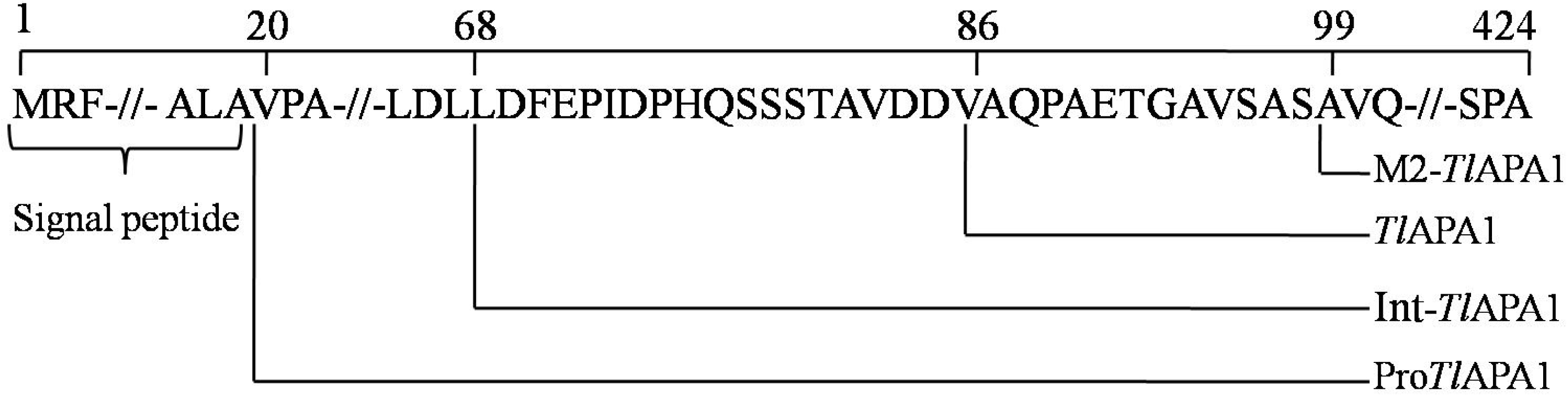
Schematic representation of the signal peptide, the propeptide region and the mature protein in *Tl*APA1. Pro*Tl*APA1, the recombinant precursor without the signal peptide; Int-*Tl*APA1, the intermediate produced during auto-activation processing; *Tl*APA1, mature protein after auto-activation at pH 3.0; M2-*Tl*APA1, mature protein after auto-activation at pH 5.0.

Generally, aspartic proteases exhibit activities and stabilities in the acidic pH range (20). Purified *Tl*APA1 showed a peak activity at pH 3.5, and was stable over the pH range of 3.0 to 6.0. These characteristics were similar to the homologs from other fungi, including *Monascus pilosus* (42), *Trichoderma asperellum* (14), and *A. foetidus* (13). The acidic adaptation of *Tl*APA1 makes it a promising candidate in many industries including cheese manufacturing, juice clarification, and leather softening. *Tl*APA1 also hydrolyzed proteins at higher temperatures. Its optimal activity was recorded at 60 °C and greater than 80% of the maximal activity was remained at 65 °C. The temperature optimum of *Tl*APA1 is higher than most reported homologs (Table 2). *Tl*APA1 retains greater than 80 % of its original activity after incubation at 55 °C for 30 min, indicating superior thermostability of *Tl*APA1 compared to most other aspartic proteases that are commonly stable at 45°C and below (Table 2). The thermostability of *Tl*APA1 is a favorable characteristic its potential application in many areas. For example, during proteolysis, most substrate proteins are resistant to proteases because of their structural stability at moderate temperatures. At higher temperatures, substrate proteins become unfold and their cleavage sites become exposed and accessible to the catalytic enzyme. This suggests more efficient hydrolysis of *Tl*APA1 substrates at high temperatures, due to the thermostable nature of *Tl*APA1.

In summary, a novel aspartic protease, *Tl*APA1, was identified, that belonged to a new clade in the phylogenetic tree. The sequence analysis of the propeptide region showed that pro*Tl*APA1 could have a unique mechanism of auto-activation processing. The results indicated that there are two steps in the processing of auto-activation, and a 45 kDa intermediate was confirmed. Moreover, the characteristics of mature protein *Tl*APA1 demonstrated that it is excellent in terms of specific activity and thermostability.

## MATERIALS AND METHODS

### Strains, vectors and substrates

The gene donor strain of *Talaromyces leycettanus* JCM12802 was purchased from Japan Collection of Microorganisms RIKEN BioResource Center. *Escherichia coli* Trans1-T1 (TransGen) was used for gene cloning and sequencing. Target gene was expressed in *P. pastoris* GS115 (Invitrogen). Cloning and expression vectors used were pEASy-T3 (TransGen, Beijing, China) and pPIC9 (Invitrogen, Carlsbad, CA), respectively. Casein sodium salt from bovine milk (C8654, Sigma-Aldrich, St. Louis, MO) was used as a substrate, and other chemicals of analytical grade were commercially available.

### Cloning of aspartic protease *Tlapa1* gene

*Talaromyces leycettanus* was cultivated as described previously (36). DNA and total RNA were extracted from the mycelia of *T. leycettanus* JCM12802 after 3 day of growth at 42 °C, and the cDNA was prepared according to the manufacturer’s instructions (TOYOBO, Osaka, Japan). The *Tlapa1* gene was amplified from DNA and cDNA of *Talaromyces leycettanus*, respectively, by polymerase chain reaction (PCR) method. The primer pairs used in this study are listed in Table 1. Finally, the PCR products were cloned into the pEASY-T3 and sequenced.

**Table 2.**
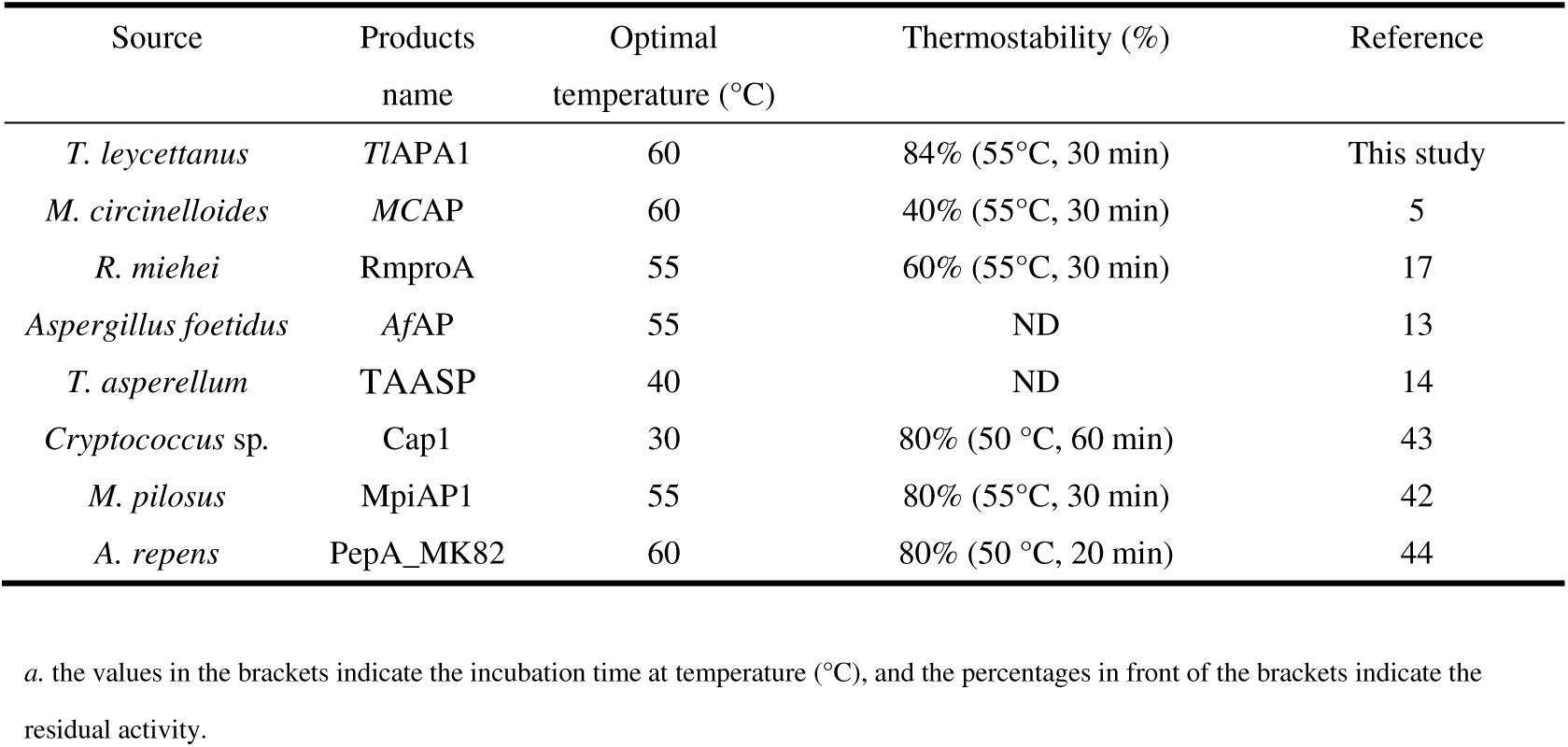
Optimal temperature and thermostability of several aspartic proteases

**Table 1.**
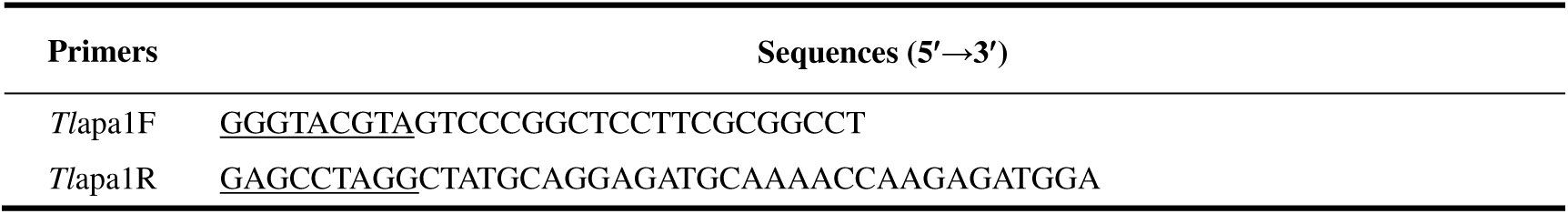
Primers used in this study.

### Bioinformatic analysis of *Tlapa1* gene

The sequence results were assembled using DNA Star 7.1 software. The amino acid sequences obtained by the Vector NTI Advance 10.0 software (Invitrogen) were searched with BLASTp programs (http://www.ncbi.nlm.nih.gov/BLAST/) to analyze the homologous sequences. The signal peptide sequence of *Tl*APA1 was predicted with SignalP (http://www.cbs.dtu.dk/services/SignalP/). The potential *N*-glycosylation sites were predicted using NetNGlyc 1.0 Server (http://www.cbs.dtu.dk/services/NetNGlyc/). Alignment of multiple protein sequences was accomplished using Clustal W software (http://www.clustal.org/) and rendered using the ESPript 3.0 program (http://espript.ibcp.fr/ESPript/cgi-bin/ESPriptcgi).

### Phylogenetic analysis

The full amino acid sequence of *Tl*APA1 was used as the query sequence in BLASTp searches in NCBI (http://blast.ncbi.nlm.nih.gov/Blast.cgi). In total, 22 sequences of A1 family aspartic proteases were obtained. Multiple sequence alignments of *Tl*APA1 with other representative aspartic proteases enzymes, characterized enzymes, and enzymes with determined three-dimensional (3D) structures, were performed as described previously (37). Sequence information for the A1 family of aspartic proteases was obtained from the MEROPS database (https://www.ebi.ac.uk/merops/cgi-bin/family_index?type=P#A). Phylogenetic analyses of *Tl*APA1 and A1 family of aspartic proteases were performed as described in previous studies (37). The distance matrix for nucleotides was calculated by Kimura’s two-parameter model. The phylogenetic tree was constructed with the neighbor-joining method using MEGA 7.0 and assessed using 1,000 bootstrap replications (46).

### Expression and purification of zymogens

Recombinant proteins were expressed in *P. pastoris* GS115, as described previously (36). Briefly, the gene fragment coding for the zymogen (pro*Tl*APA1) without the signal peptide was amplified using PCR method. PCR products were digested with *Eco*RI and *Not*I and ligated into the pPIC9 plasmid using T4 DNA ligase (New England Laboratory). The recombinant plasmid pPIC9-pro*Tlapa1*, linearized by *Bgl*II, was transformed into *P. pastoris* GS115 competent cells by electroporation. Positive transformants were screened based on the transparent zone on skim milk plates as described below. The transformants showing the largest transparent zones were inoculated into 30 mL YPD and incubated at 30 °C. The seed medium containing the positive transformant was inoculated into 1 L conical flasks containing 300 mL of BMGY for fermentation. Conical flasks containing 200 mL of BMMY and 0.5% (v/v) methanol were prepared.

The cells were harvested by centrifugation for 10 min at 12,000*g*, and resuspended in BMMY medium and for next subsequent fermentation at 30 °C. Methanol was added every 24 h to obtain a final concentration of 0.5% (v/v). After 48h of cultivation, cell-free cultures were centrifugated at 12,000 rpm, 4 °C for 10 min and fermentation broth was collected. The crude pro-enzymes were concentrated using an ultrafiltration membrane with a molecular weight cut-off of 10 kDa (Vivascience, Hannover, Germany). A HiTrap Q Sepharose XL 5 mL FPLC column (GE Healthcare, Sweden) was used for purification. Protein binding and equilibration was performed using buffer A (10 mM sodium phosphate, pH 6.0), and a linear gradient of NaCl (0–1.0 M) was used to elute the proteins.

### Activation of purified zymogen pro*Tl*APA1

To determine processing of zymogen conversion, pH of the pro*Tl*APA1 samples were adjusted to 3.0 using 0.5 M lactic acid-sodium lactate buffer. pro*Tl*APA1 was auto-catalytically activated at 37 °C for 0, 15, 30, 45, 60, 75, and 90 min, respectively. Processed polypeptides were detected by SDS-PAGE and N-terminal sequencing. To study the effect of pepstatin A on the zymogen conversion of pro*Tl*APA1, 5 μM pepstatin A was added into the conversion system before he auto-catalytic processing and incubated at 37 °C for 15, 30, 45, and 60 min, respectively. Zymogen conversion systems were subjected to sodium dodecyl sulfate-polyacrylamide gel electrophoresis (SDS-PAGE) analysis. Samples without pepstatin A were also treated similarly, as a control group. To determine the optimum pH of zymogen conversion, the pH of the pro*Tl*APA1 samples were adjusted to 2.0, 3.0, 4.0, 5.0, and 6.0 using 0.5 M lactic acid-sodium lactate buffer. The samples at different pH conditions were incubated at 37 °C, and the proteolytic activities were analyzed with casein (1%, w/v) at different incubation times. Finally, the samples were analyzed using SDS-PAGE.

### Enzyme activity assay

The activity of aspartic proteases was assayed in a 1000 μl reaction mixture containing 500 μl of 1 % (w/v) casein sodium salt and 500 μl enzyme sample in buffer at pH 3.0. After incubation at 60 °C for 10 min, 1000 μl of 40 % (w/v) trichloroacetic acid (TCA) was added to terminate the reaction. 500 μL of the supernatant was obtained from the mixture using centrifugation after 12,000*g* for 3 min. 2.5 mL of 0.4 M sodium carbonate and 500 μL Folin-phenol was added into supernatant in turn before incubating at 40 °C for 20 min. The amount of released tyrosine was measured at 680 nm. One unit of proteolytic activity was defined as the amount of enzyme that released 1 μmol of tyrosine equivalent per minute under the conditions described above (pH 3.5, 60 °C and 10 min).

### Properties of recombinant aspartic protease *Tl*APA1

Optimal conditions for purified *Tl*APA1 activity were measured in lactic acid-sodium lactate buffer under the following conditions– temperatures ranging from 30 °C to 80 °C (at a constant pH 3.5); and pH ranging from 2.0 to 4.5 (at a constant temperature of 60 °C). Thermostability and pH stability of *Tl*APA1 were assessed by preincubating the purified enzyme for 1 h in lactic acid-sodium lactate buffer under varying temperature conditions: 55 °C, 60 °C and 65 °C (at constant pH of 3.5); or varying pH conditions: pH 2.0–11.0 (at constant temperature of 37 °C), respectively, and then determining the residual enzyme activity.

The proteolytic activities of *Tl*APA1 were measured under standard conditions (pH 3.5, 60 °C, 10 min) with 0.5–10 mg/mL casein sodium salt, and the constants were determined by linear regression fitting using GraphPad Prism version 7.01. All experiments were performed in three biological and technical replicates.

### N-terminal Sequencing

Purified proteins were separated by SDS-PAGE and electro transfered onto a polyvinylidene difluoride (PVDF) membrane. Stained with Coomassie Brilliant Blue R-250, the target protein bands were excised and subjected to N-terminal amino acid sequence analysis using a PPSQ-33 automatic sequence analysis system (Shimadzu, Kyoto, Japan).

## ACKNOWLEDGEMENTS

This work was supported by the National Key Research and Development Program of China (2016YFD0501409-02), the Fundamental Research Funds for Central Non-profit Scientific Institution (Y2017JC31), and the China Modern Agriculture Research System (CARS-41).

